# BRCA1 expression, its correlation with clinicopathological features and response to neoadjuvant chemotherapy in high grade serous ovarian cancer from an Indian centre

**DOI:** 10.1101/2023.01.11.523693

**Authors:** Akkamahadevi Patil, Sharada Patil, CE Anupama, Savitha Rajarajan, Vidya Prasad Nimbalkar, Usha Amirtham, G Champaka, MN Suma, Geetha V Patil, Ashwini Nargund, VR Pallavi, Linu Jacob, CS Premalatha, Jyothi S Prabhu

## Abstract

In high grade serous ovarian cancers (HG-SOC), BRCA1/2 mutations have been reported as the most predominant mutations by various studies. However, the non-mutational mechanisms of BRCA pathway inactivation in HG-SOC are unclear. We aimed to evaluate BRCA1 inactivation by estimating its expression along with its repressor in primary and neoadjuvant chemotherapy (NACT) treated HG-SOC tumors with known therapeutic response.

The expression pattern of BRCA1 protein was evaluated by immunohistochemistry (IHC) in 119 cases of HG-SOC from a hospital cohort consisting of primary (N= 69) and NACT treated (N=50) tumors. Histological patterns (SET), stromal infiltration by lymphocytes (sTILs) and chemotherapy response score (CRS) were estimated by microscopic examination. Gene expression levels of BRCA1, and its repressor ID4 was estimated by qPCR. Association of BRCA1 protein and mRNA with clinicopathological features was studied. Relevance of the BRCA1/ID4 ratio was evaluated in tumors with different CRS.

BRCA1 protein expression was observed in 12% of primary and 19% of NACT treated HG-SOC tumors. Moderate concordance was observed between BRCA1 protein and mRNA expression (AUC-0.677). High BRCA1 mRNA expression was significantly associated with more frequent SET pattern (p=0.024), higher sTILs density (p=0.042), increased mitosis (p=0.028). BRCA1 negative tumors showed higher expression of ID4 though not statistically significant. Higher BRCA1/ID4 ratio was associated with high sTILs density in primary (p=0.042) and NACT treated tumors (p=0.040). Our findings show the utility of BRCA1/ID4 ratio to predict neoadjuvant therapy response, which needs further evaluation in larger cohort with long term outcomes.

## INTRODUCTION

Epithelial ovarian cancer (EOC) is a complex disease. It is the second most common cause of death among the gynecological cancers, accounting for 2.5% of all malignancies among females but contributes to 5% of the female cancer deaths. This disproportionate low survival rates are due to the diagnosis in the late stage ^1^. The current standard treatment for EOC is cyto-reductive surgery and adjuvant platinum-based chemotherapy. Neoadjuvant chemotherapy is considered for patients with extensive disease and having difficulty for complete cytoreduction. Despite standard treatment recurrence is very common in EOC and these recurrent tumors are not responsive to chemotherapy due to the development of the chemoresistance ^2^.

Among five different subtypes of EOC, High Grade Serous Ovarian Cancers (HG-SOC) is the dominant subtype diagnosed clinically, constitute majority (70%) of ovarian tumors and has poor 5 years survival rates of 5-30% despite the standard treatment. HG-SOC is characterized by genomic instability with deficiency in DNA repair pathways such as DNA double strand break repair by homologous recombination. BRCA1/2 and many others are involved in these repair pathways ^3^. BRCA1/2 dysfunction leads to homologous recombination deficiency (HRD), due to which the cells are unable to utilize the homologous recombination repair pathway when double strand breaks occur ^4^. Women with germline mutations in BRCA1 or BRCA2 have a 30% to 70% chance of developing ovarian cancer by the age of 70, with most cases associated with HG-SOC ^5^. BRCA pathway dysfunction seems to be a hallmark of HG-SOC. BRCA dysfunction is known to occur through multiple mechanisms like mutation, activation of trans acting repressors, epigenetic mechanisms like DNA methylation and miRNA mediated suppression. Several studies have shown the better response to commonly used DNA damaging drugs like platinum salts in ovarian cancer in patients with dysfunction of BRCA1 pathway.

Substantial progress in the past decade in the understanding of genetic and molecular mechanism of HG-SOC tumor has shown BRCA1/2 as one of the predominant mutations, but the non-mutational mechanisms and the overall frequency of BRCA pathway inactivation in HG-SOC is unclear at present. We evaluated BRCA1 inactivation by estimating its expression along with its repressor in tumor samples of primary and neoadjuvant chemotherapy (NACT) treated HG-SOC with known therapeutic response.

## METHODS AND MATERIALS

### a) Cohort of HG-SOC cases

In the current study, BRCA1 dysfunction was evaluated by estimating its protein and mRNA expression and correlating it with its repressors in tumor samples of HG-SOC with known chemotherapeutic response. The study included 119 cases that were retrieved from the department of Pathology, Kidwai Memorial Institute of Oncology (KMIO) which were diagnosed with HG-SOC and operated between the year 2016 to 2019. Ethical approval was obtained for the study from the institutional ethics committee (KCI/MEC/027/10.Agust.2018). Tumor samples having ≥50% tumor content was considered. Among these 119 tumor samples, 69 were primary tumor samples and 50 were post neoadjuvant chemotherapy (NACT) tumor samples. Of these post NACT samples (n=50), chemotherapy response score (CRS) was available for 42 cases.

Microscopic evaluation of histological features: All the tumor sections of HG-SOC cases were evaluated under the scanner view of the microscope (4x) for various architectural types such as solid, pseudoendometrioid (glandular) and transitional cell carcinoma like patterns (SET pattern) as these features are commonly expressed in BRCA1 mutation. The percentage of these SET pattern in each tumor was estimated after screening all the tumor sections. Other architectural features such as papillary, micropapillary and infiltrative papillary were documented. Presence of tumor infiltrating lymphocytes (TILs) was evaluated in the hotspots disregarding the lymphocyte aggregates at the periphery of tumor nests. Similarly, mitotic index was calculated by counting mitosis per 10 high power fields, in hotspots. Proportion of necrosis was identified by screening all the tumor sections and documented as percentage of the total area.

Chemotherapy response scores (CRS) was evaluated according to CAP guidelines ^6^ for histopathologic assessment of response to neoadjuvant chemotherapy. Briefly in the three-tier scoring system CRS1 indicated no or minimal tumor response, CRS2 was appreciable tumor response amidst viable tumor, CRS3 was with complete or near complete response with no residual tumor.

### b) Immunohistochemistry (IHC)

IHC for the BRCA1 protein was performed as detailed in the previous study ^7^. Briefly, primary antibody for BRCA1 (Clone MS110, Calbiochem Cat# OP92, Darmstadt, Germany) was used at 1:100 dilution and incubated for 1 hour at room temperature. After this, sections were incubated with secondary antibody (DAKO REAL™ EnVision ™ Glostrup, Denmark) for 20 min as per the kit instructions, followed by colour development using DAB (DAKO REAL™ EnVision ™) as chromogenic substrate for 10 minutes. Sections were counterstained with hematoxylin and mounted after dehydration in graded alcohol and xylene. Appropriate positive and negative controls were run for each batch.

### c) Extraction of Nucleic acids from the selected FFPE tumor samples

Total ribonucleic acid (RNA) was extracted from two 20μm thick sections from each patient’s tumor tissue block and these sections were deparaffinized by heat, followed by overnight digestion using proteinase K (Qiagen #19133). Total RNA was then extracted using Tri Reagent protocol according to the manufacturer instructions (Sigma Aldrich # T9424). RNA Quantitation was done using the Ribogreen dye (Invitrogen # R11490-Quant –iT Ribogreen RNA assay kit) on a fluorescent plate reader (Tecan M200-Pro Infinite Series). A total of 500ng RNA was reverse transcribed to cDNA using the ABI high-capacity cDNA archive kit (ABI # 4322171) as per the manufacturer’s protocol.

### d) Gene expression of BRCA1 and ID4 and normalization

Expression level of test genes (BRCA1 and ID4) along with a panel of 3 reference genes (PUM1, RPL13A, ACTB) was performed using TaqMan qPCR chemistry on the Light Cycler 480 II (Roche Diagnostics). According to previously established methods ^7^. To determine relative transcript abundance in these samples, the Ct values for the test gene were normalized to the mean Ct value of the three reference genes (house-keeping genes) for each sample as ΔCt. The Relative Normalized Units (RNU) of expression of the test genes was calculated as 15-ΔCt.

## RESULTS

The median age of the cohort was 52 years and the majority (75%) of the cases were bilateral. SET pattern was observed in half the cases, high TILs density was observed in more than 50% of primary tumors. Clinical features are represented in Table 1.

**Table 1.** Clinicopathological features of the study cohort.

| Variables |  | N=119 (%) |
| --- | --- | --- |
| Age (years) | Mean | 52.3 |
|  | Median | 52 |
| Laterality | Unilateral | 24 (20) |
|  | Bilateral | 73 (62) |
|  | Not mentioned | 22 (18) |
| Specimen type | Primary | 69 (58) |
|  | Post NACT | 50 (42) |
| SET pattern | Present | 60 (51) |
|  | Absent | 59 (50) |
| Necrosis | Present | 64 (54) |
|  | Absent | 54 (45) |
|  | Not mentioned | 01 (1) |
| sTILs | High TILs | 38 (32) |
|  | Low TILs | 30 (25) |
|  | Not mentioned | 51 (43) |
| Mitosis/ HPF | High mitosis | 32 (27) |
|  | Low Mitosis | 36 (30) |
|  | Not mentioned | 51 (43) |
| BRCA1 IHC | Positive | 17 (15) |
|  | Negative | 98 (82) |
|  | Not mentioned | 04 (3) |

sTILs- stromal tumor infiltrating lymphocytes; HPF-High power field; IHC-Immunohistochemistry

### BRCA1 protein expression

BRCA1 IHC was performed on all 119 cases. Among these, IHC staining could not be assessed in four cases due to floating of the tissue sections. Nuclear and/or cytoplasmic staining was observed in tumor cells. 10% and above of the tumor cells stained was considered positive.

Out of 115 cases assessed, 17 cases (14.7%) were positive for BRCA1 staining which showed the protein in the nucleus (with or without in the cytoplasm) and 98 (85.3%) were negative. Representative BRCA1 IHC images are shown below figure 1 (A and B). 8/68 (11.8%) primary treatment naïve tumors were BRCA1 positive and 9/47 (19.2%) NACT treated tumors showed BRCA1 positivity.

**Figure 1.**
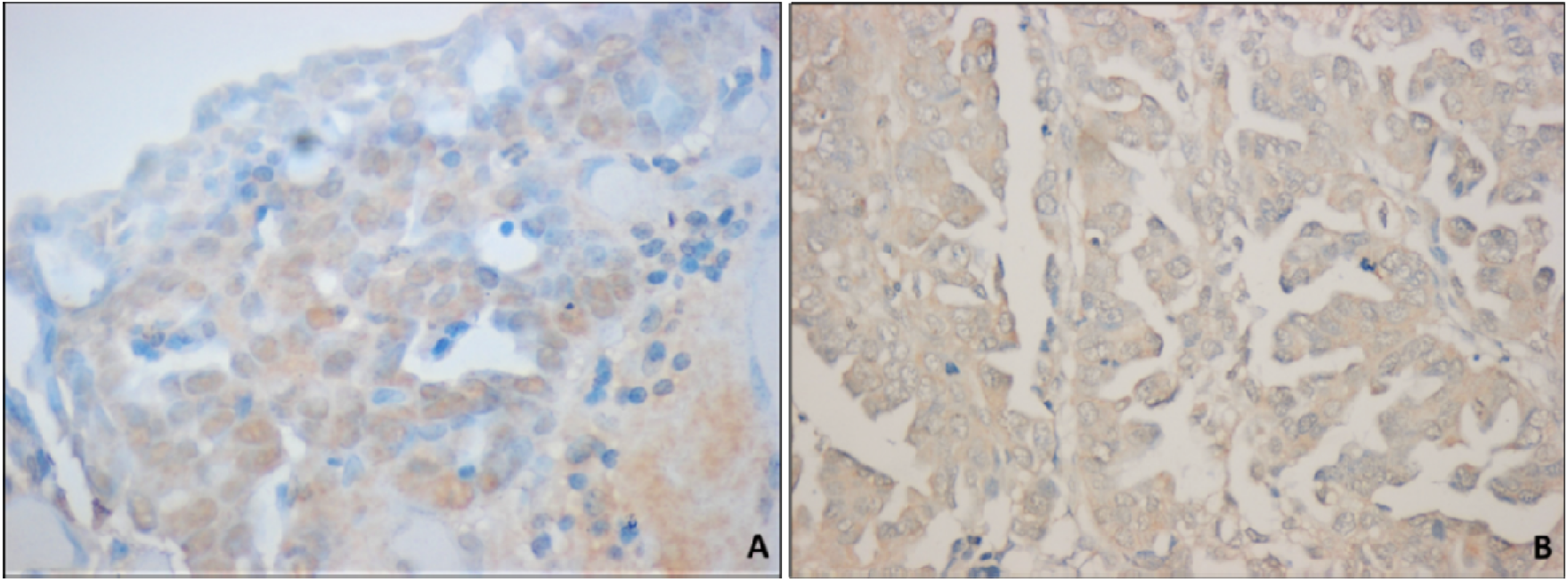
BRCA1 immunohistochemistry Images. A. Nuclear staining 20x. B. Cytoplasmic staining 20x

### Expression and distribution of BRCA1 and ID4 expression in the primary and NACT treated tumors

BRCA1 and ID4 expressions were assessed in 108 cases out of 119 cases based on quality and quantity of RNA available. It was observed that BRCA1 mRNA [median(range)] expression was significantly higher (p=0.009) in primary tumors (n=66) [RNU=8.24(1.45-11.81)] compared to NACT treated tumors (n=42) [RNU=7.46 (4.05-10.32)]. However, we did not find any significant difference (p=0.411) in the expression of ID4 in primary [12.583 (9.58-15.76)] and NACT treated [12.278(9.43-16.09)] tumors.

### Association of BRCA1 with clinical features in primary tumors

Association of BRCA1 protein expression with various clinical features, such as age, laterality, SET pattern, necrosis, TILs density and mitosis were assessed in primary tumors. As the number of BRCA1 positive cases was low, we did not observe any statistically significant correlation with the assessed clinical parameters.

Next, we evaluated the concordance between BRCA1 protein expression and its transcript, using receiver operating characteristic (ROC) curve analysis. Analysis showed moderate concordance between protein and transcript with area under curve (AUC=0.677) showing trends towards the significance (p=0.077). As BRCA1 transcript showed moderate concordance with the protein expression, association of BRCA1 mRNA expression with different clinical features was evaluated. BRCA1 mRNA expression in primary tumors ranged from RNU 1.45-11.8. Based on the third quartile value (RNU= 9.60) primary tumor samples were divided into BRCA1 high and BRCA1 low groups. Association of clinical features such as SET pattern, necrosis, TILs and mitosis with BRCA1 high and low groups was examined.

Higher BRCA1 mRNA expression was associated with more frequent SET pattern (p=0.024), higher stromal TILs density (p=0.042), increased mitotic indices (p=0.028). BRCA1 mRNA expression did not show any association with necrosis (p=0.805) as mentioned in table 2.

**Table 2.**
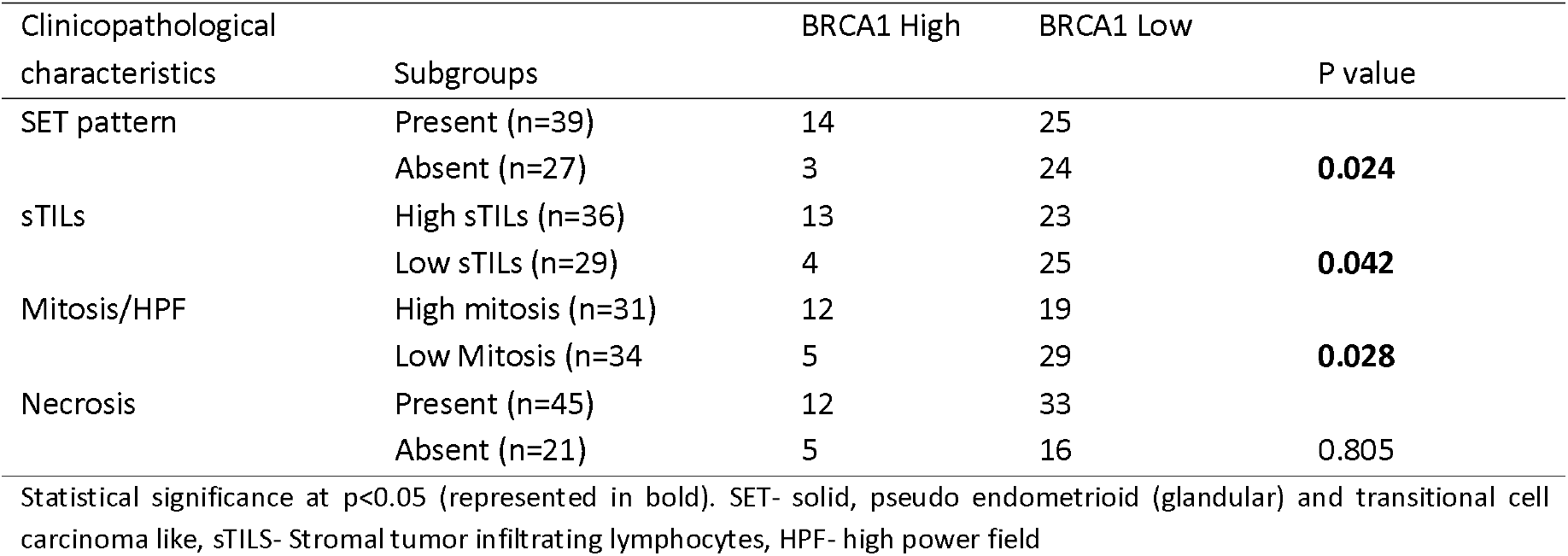
Association of BRCA1 mRNA expression with clinicopathological features in primary HG-SOC.

### Relationship between BRCA1 and its repressor ID4

We evaluated correlation of BRCA1 and ID4 transcripts in BRCA1 positive tumors (based on IHC) (n=7). On analysis, it was observed that ID4 expression was negatively correlated with BRCA1 transcript (r= - 0.321), though it was not statistically significant (p=0.498). This might be due to the smaller number of cases that were BRCA1 positive by IHC. We also examined ID4 expression in BRCA1 positive and negative tumors. It was observed that BRCA1 negative tumors showed higher ID4 expression compared to BRCA1 positive tumors, though the difference was not statistically significant (p=0.246).

### Clinical significance of BRCA1/ID4 ratio in primary and NACT treated tumors

To understand the clinical significance of cumulative effect of BRCA1 and ID4, BRCA1/ID4 ratio was calculated. Ratio ranged from 0.127-1.005 with a mean value of 0.641. Based on third quartile value (0.741), BRCA1/ID4 ratio was grouped in high and low groups.

Association of BRCA1/ID4 high and low groups with clinical parameters showed that the tumors with high BRCA1/ID4 ratio had high stromal TILs density (p=0.042). There was no association of BRCA1/ID4 ratio with other clinicopathological parameters.

Among post NACT treated tumors chemotherapy response score (CRS) and BRCA1/ID4 ratio was available for 36 tumors. These tumors were grouped into two groups based on the response as CRS1 (No response, n=21) and CRS2 (with partial response, n=15). It was observed that non-responsive tumors had increased BRCA1/ID4 ratio (p=0.141) as shown in figure 2.

**Figure 2.**
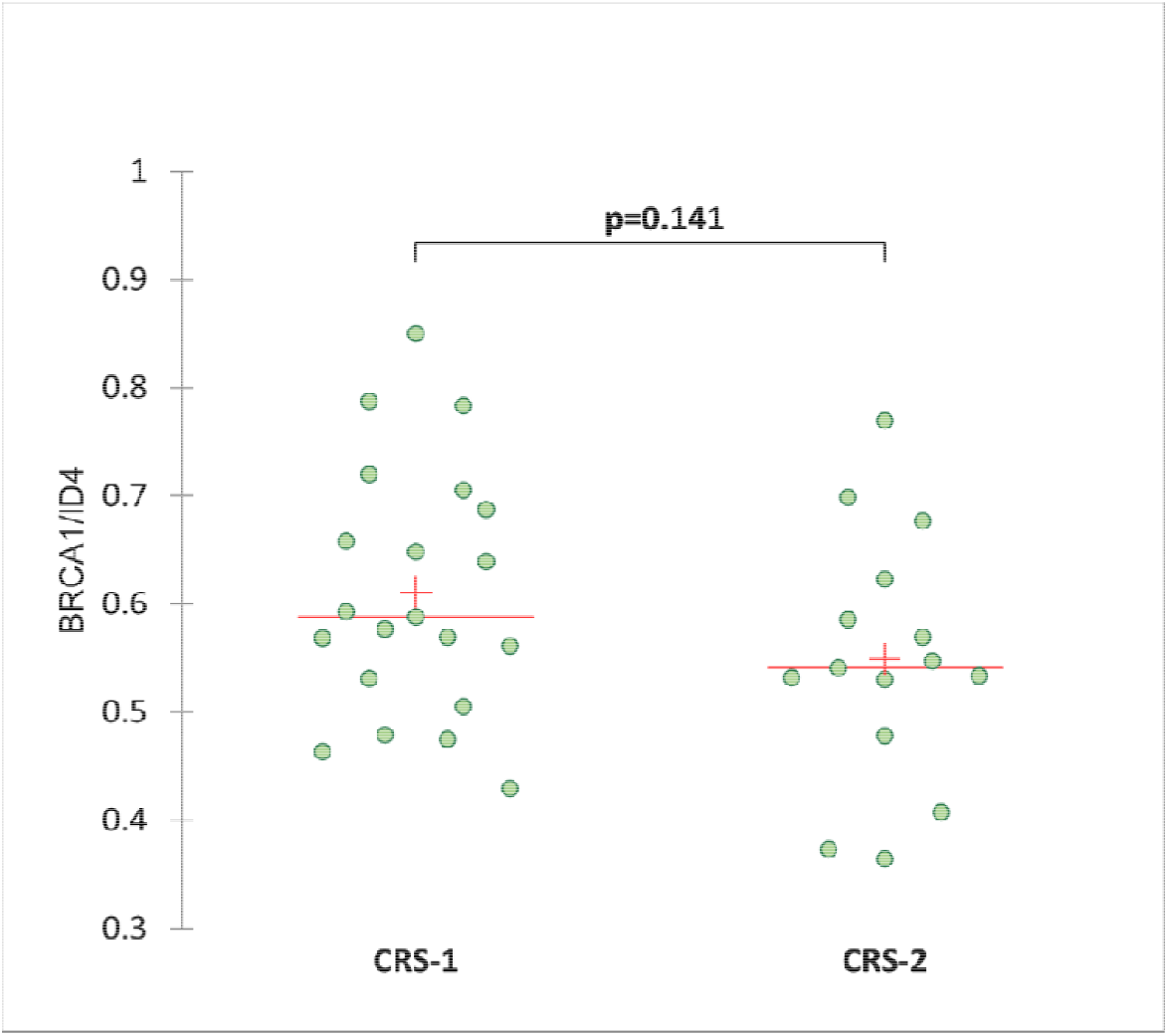
Association of BRCA1/ID4 ratio with chemotherapy response score (CRS)

In line with the primary tumors, BRCA1/ID4 high ratio was associated with high stromal TILs density in NACT treated HG-SOC tumors (p=0.040). Additionally, we also observed that in NACT treated tumors, CRS-1 response was associated with necrosis (p=0.010). There was no association of CRS score with stromal TILs density.

## DISCUSSION

HG-SOC is the most common subtype of ovarian cancer. It is mostly diagnosed at the advanced stages (stage III and IV) and accounts for 70-80% of the deaths from all forms of ovarian cancer. Despite the extensive research, the overall survival has not improved for decades ^8^. In 2011 TCGA study explored the genomic nature of ovarian cancer. This study found 96% of HG-SOC patient samples had TP53 mutations suggesting that it was the key alteration involved in the disease initiation. Beside this the study also reported that BRCA1 mutation in 12.1% and BRCA2 mutation in 11.5% HG-SOC cases which included both germline and somatic BRCA mutations. Further pathway analysis showed homologous recombination DNA repair (HR) pathway was defective in 51% HG-SOC cas?es. Among the patients with defective pathway 20% had BRCA mutations and 11% had BRCA silencing by DNA hypermethylation ^9^. A study by Sun et al. assessed miR9 expression in HG-SOC and demonstrated that its high expression down regulates BRCA1 expression making it more sensitive to the platinum based therapy and thus improves the progression free survival ^10^. Many others studies have reported that BRCA dysfunction due to various mechanisms are associated with better survival in HG-SOC patients as they respond better to platinum-based chemotherapy compared to BRCA wildtype HG-SOC ^9,11^. However, there are limited studies that have evaluated the clinical significance of BRCA1/2 mRNA and protein expression in HG-SOC its association with the therapy response.

In this study we assessed the association of BRCA1 expression with various morphological features of HG-SOC. Though we did not find any association of BRCA1 protein expression with these morphological features, high BRCA1 mRNA expression was associated with more frequent SET patterns, high stromal TILs density and higher mitotic index. Our findings are in concordance with previous study reporting BRCA1 association with SET features, higher mitotic indexes, more TILs and either geographic or comedo necrosis in HG-SOC ^6^. In our study we did not find an association of BRCA expression with necrosis in primary tumors. However, in NACT treated tumors high BRCA1/ID4 ratio was associated with poor chemotherapy response. These chemoresistant tumors were associated with presence of necrosis.

ID4 is an important dominant negative transcriptional repressor gene involved in regulating BRCA1 expression and might therefore be important for the BRCA1 regulatory pathway involved in the pathogenesis of sporadic breast and ovarian cancer ^12^. Hence, we calculated the BRCA1/ID4 ratio to understand the effect of BRCA ness in HG-SOC. We noticed inverse trends in BRCA1 and ID4 expression BRCA1 positive tumors and BRCA1 negative tumors showed comparatively increased ID4 expression. These results suggest that ID4 represses BRCA1 gene expression in addition to other repressor/regulators involved in BRCA 1 dysfunction.

In our study, the increased BRCA1/ ID4 ratio was associated with increased stromal TILS density in both primary and NACT treated HG-SOCs. TILs are involved in elucidating the antitumor immune response. Various studies have assessed the association between TILs and prognosis in HG-SOC patients ^13–15^. However, results across different studies are conflicting. A recent study has demonstrated that BRCA1/2 mutated HG-SOCs harbor increased CD3+ and CD8+ TILs and elevated PD-1 and PD-L1 expression compared to HR proficient HG-SOCs. Survival analysis showed that both BRCA1/2-mutation status and number of TILs were independently associated with outcome. Within HGSOCs, a group with HR proficient (BRCA wild type) and low number of TILs were associated with very poor prognosis and the other group with BRCA1/2-mutated tumors and high number of TILs was associated with very good prognosis HG-SOCs ^16^. These findings suggest that BRCA-mutated HG-SOCs harbor anti-tumorigenic TILs and these tumors might be more sensitive to PD-1/PD-L1 inhibitors compared to BRCA wildtype tumors.

In NACT treated tumors high BRCA1/ID4 ratio was associated with poor response to platinum-based therapy in our study. This was in line with the previous study by Tsibulak et al. demonstrating that BRCA mutant tumors express low BRCA mRNA and these tumors had higher sensitivity to platinum-based therapy ^17^. Many other studies have reported that BRCA mutant subtype of HG-SOC, which are homologous recombination repair deficient respond better to the platinum-based therapy ^18,19^. On the other hand, BRCA wildtype or homologous recombination repair proficient subtype of HG-SOC are more likely resistant to platinum therapy.

We observed in primary tumors, BRCA1 expression is associated with characteristic morphological features of HG-SOC. This association was not observed in NACT treated tumors. In both primary and NACT treated tumors high BRCA1/ID4 expression was associated with increased stromal TILS density. The novelty of our study lies in using simple gene expression approach to understand the association of BRCA1/ID4 expression with various clinicopathological features and predicting the therapy response in HG-SOC. However, this study has significant limitations. As it is a retrospective study, all clinical features were derived from medical records. Given the problems associated with detection of BRCA1 protein by IHC using the antibody clone (MS110), very low proportion of tumors showed BRCA1 protein expression and the association with many parameters did not reach statistical significance despite showing the expected trends. Because of the lack of long term follow up and prognostic data in the series, we were unable to derive prognostic significance of the measured parameters.

## CONCLUSION

Findings of our study support the evidence of low expression of BRCA1 in HG-SOC. Apart from mutational mechanisms BRCA1 dysfunction could be due to higher expression of its repressors such as ID4 and evaluation of BRCA1/ID4 ratio might be helpful to identify therapy resistant tumors. Our findings should be evaluated in larger cohorts with long term outcomes to identify prognostic and therapeutic significance.

## Acknowledgements

The authors would like to acknowledge Rajiv Gandhi University of Health Sciences (RGUHS) for funding the study.

